# Waves of Maturation and Senescence in Micro-Structural MRI Markers of Human Cortical Myelination over the Lifespan

**DOI:** 10.1101/314195

**Authors:** Håkon Grydeland, Petra E. Vértes, František Váša, Rafael Romero-Garcia, Kirstie Whitaker, Aaron F. Alexander-Bloch, Atle Bjørnerud, Ameera X. Patel, Donatas Sedervicius, Christian K. Tamnes, Lars T. Westlye, Simon R. White, Kristine B. Walhovd, Anders M. Fjell, Edward T. Bullmore

## Abstract

Seminal human brain histology work has demonstrated developmental waves of myelination. Here, using a micro-structural magnetic resonance imaging (MRI) marker linked to myelin, we studied fine-grained age differences to deduce waves of growth, stability, and decline of cortical myelination over the life-cycle. In 484 participants, aged 8-85 years, we fitted smooth growth curves to T1- to T2-weighted ratio in each of 360 regions from one of 7 cytoarchitectonic classes. From the first derivatives of these generally inverted-U trajectories, we defined three milestones: the age at peak growth; the age at onset of a stable plateau; and the age at the onset of decline. Age at peak growth had a bimodal distribution comprising an early (pre-pubertal) wave of primary sensory and motor cortices and a later (post-pubertal) wave of association, insular and limbic cortices. Most regions reached stability in the 30s but there was a second wave reaching stability in the 50s. Age at onset of decline was also bimodal: in some right hemisphere regions, the curve declined from the 60s, but in other left hemisphere regions, there was no significant decline from the stable plateau. These results are consistent with regionally heterogeneous waves of intracortical myelinogenesis and age-related demyelination.

## Introduction

The cortex contains myelinated axons (Gennari 1782) critical for efficient neuronal communication (Zalc and Colman 2000). More than a century ago, Paul Flechsig (1847-1929) noted that cortical myelination did not occur simultaneously in all regions, but rather in a succession of waves peaking at different ages (Flechsig 1901). Motor and sensory cortical regions myelinated earlier than association cortical regions. Yakovlev and Lecours (1967)extended this notion of waves of myelinogenesis, and noted how intracortical fibres of the association cortex appeared to have a long myelination cycle, demonstrating an immature “greyish pallor” beyond the third decade of life. Convergently, studies in rodents have shown myelin-forming oligodendrocyte cells are produced in waves (Kessaris et al. 2006). Specifically, three waves of oligodendroglial proliferation have been described, peaking at different stages of forebrain development before and after birth (Rowitch and Kriegstein 2010). Later in life, primate histology studies have reported cortical myelin degeneration, and aberrant remyelination (Peters 2009).

Magnetic resonance imaging (MRI) has allowed for the indirect study of myelin using volume and other measures likely related to myelin in development (Paus et al. 1999), adulthood, and aging (Bartzokis et al. 2001). Such studies, focusing mainly on the white matter (WM) of the cerebrum, seem to largely support Flechsig’s idea of myelination heterochronicity (Gao et al. 2009). van der Knaap and Valk (1990)studied myelination during the first postnatal year using T1- and T2-weighted images, delineating 5 stages, or waves, of myelination. However, there have been no attempts to test Flechsig’s notion of cortical waves during childhood and adolescence, and whether similar patterns could be found late in life.

There have been only a few studies encompassing a sufficient age range to investigate putative myelinogenesis, from childhood to old age, comprehensively across the human cortex. Studying the WM, Kochunov et al. (2012)fitted linear models to diffusion tensor imaging data before and after the age at peak fractional anisotropy (a marker of WM tract organization) in participants aged 11-90 years. In nine WM tracts, they reported that tracts with a faster rate of maturation had a faster rate of age-related decline. Yeatman et al. (2014)fitted a piecewise linear model of R1 in 24 WM pathways for participants aged 7-85 years. R1, a measure of the longitudinal relaxation rate of protons excited in the imaging process (Sigalovsky et al. 2006), is a quantitative marker sensitive to myelin. They found no systematic relationship between the age at which a tract matures and the age at which it begins to decline. Arshad et al. (2016)studied 6 WM tracts in 61 adults aged 18-84 using myelin water volume fraction (a quantitative measure sensitive to myelin), and reported quadratic age effects. Importantly, a few studies have looked at cortical myelination. Deoni et al. (2015)found logarithmic increases in myelin water volume fraction with age in children aged 1 to 6 years. Using methods sensitive to myelin based on magnetization transfer, Whitaker et al. (2016)showed a linear cortical myelin increase for participants aged 14-24 years. Callaghan et al. (2014)reported stable levels of myelin until around 60 years of age, followed by decrease, in several cortical regions including the Heschl’s gyri, for participants aged 19-75 years.

The lifespan studies (Kochunov *et al.* 2012; Yeatman *et al.* 2014) yielded mixed support for the concept that decline occurs in reverse order to growth of myelination. This so-called “last in, first out” hypothesis (Raz 1999) suggests that regions that myelinate last in the course of development, are the first to demyelinate later in life, i.e. are more affected by age. This heterochronic pattern has been related to association cortices compared with sensory cortices (Raz 1999), possibly due to their underlying cytoarchitecture. The association cortex is particularly well connected, specialised for multimodal integration, and underpins most complex behaviours (van den Heuvel and Sporns 2013). Therefore, these regions might need to remain plastic for as long as possible to refine their connections (Glasser et al. 2014) and consolidate them by a prolonged phase of myelination. In line with this reasoning, but in contrast to the “last in, first out” hypothesis, these regions could be the last to decline (“last in, last out”). The high connectivity of the association cortex, and the importance of myelin in brain network functioning, also raise the question of how patterns of myelination relate to how the brain is organised as a network. Particularly, whether the most well-connected “hub” regions are later to mature and to decline (Whitaker *et al.* 2016).

In this context, we aimed to investigate myelination across the cortex in humans (N=484, aged 8-85 years), using a micro-structural MRI marker putatively linked to myelin (Glasser *et al.* 2014; Glasser et al. 2016): the ratio of T1- and T2-weighted MRI images (T1w/T2w). Although not a truly quantitative measure of intracortical myelin, and likely influenced by other characteristics within each voxel (Glasser *et al.* 2016), variation in T1w/T2w has been shown to match histologically-derived myelin content (Glasser and Van Essen 2011). Studies comparing cortical T1 and T1w intensity with myelin stains in marmosets (Bock et al. 2009; Bock et al. 2011), have supported the interpretation of the T1w/T2w ratio as a reasonable estimate of relative intracortical myelin content (Glasser *et al.* 2014). Thus, while acknowledging that the T1w/T2w ratio is far from a pure measure of myelin content, we adopt this interpretation here, as in our previous work. T1w/T2w variation has been related to individual differences in electrophysiology (Grydeland et al. 2015), and cognition (Grydeland et al. 2013), and has shown positive relationships with age in development and early adulthood, and negative relationships with age in later life (Grydeland *et al.* 2013; Shafee et al. 2015). Here, we move further by testing Flechsig’s theory of waves across the lifespan, and by testing for associations between development and aging. From the T1w/T2w maps, we fitted smooth age-trajectories for 360 cortical regions (Glasser *et al.* 2016) in each of 7 cytoarchitectonic classes (Scholtens et al. 2016). The first derivatives of these cross-sectionally estimated cortical growth curves allowed us to define three key micro-structural milestones, namely age at peak growth, age at onset of stability, and age at onset of decline. We analysed the distribution of these milestones for deviations from unimodality (i.e. at least bimodality). We related the timing of the milestones both to (i) the estimated maximum rates of maturation and decline in each region; and (ii) the topological properties of each node in the structural covariance network estimated on average across a subset of participants, as well as a functional connectivity network based on functional MRI scans.

On this basis, we tested three hypotheses: i) the timing of micro-structural milestones – age at peak growth, age at onset of stability, and age at onset of decline - has at least a bimodal distribution indicative of successive waves of change in myelination, ii) timing of these milestones is related to rates of micro-structural maturation and decline, and iii) the variation in milestones and rates of micro-structural growth and decline relate to cytoarchitectonic properties, and network topology, namely connectivity strength or hubness of regions in the anatomical network. We particularly expected the association cortex to develop and decline differently compared with the most heavily myelinated primary motor and sensory cortices.

## Materials and Methods

### Participants

Participants were included after screening for conditions assumed to affect CNS function (for details regarding *Methods*, see ***Supplementary Material***). Based on the existence of a T1w and a T2w scan for each individual, and after stringent scan quality control (see below), we included 484 participants: 263 females (54.3%), mean age (SD) = 38.3 (22.5) years, median age= 34.6 years, age range = 8.2-85.4 years. ***Supplementary Figure S1*** shows the gender and mean full-scale intelligence quotient for females and males across age deciles.

### MRI Data Acquisition, Quality Control, and Processing

Scans were acquired using a 12-channel head coil on a 1.5-T Siemens Avanto scanner (Siemens Medical Solutions, Erlangen, Germany) at Oslo University Hospital Rikshospitalet, in one session per participant, using a 3D T1w magnetization-prepared rapid gradient echo (T1w), and a 3D T2w sampling perfection with application optimized contrasts using different flip angle evolutions sequence (T2w). All raw scans were visually inspected to assess quality and identify motion-related artefacts. A total of 18 participants were excluded due to incomplete T2w scans, overfolding, or movement artefacts in T1w, T2w, or both. Of these, 6 scans were excluded due to signs of motion (all males, 4 participants aged between 9 and 12 years, and 2 participants aged 52 and 68 years). In evaluating this number of motion-compromised scans, it is relevant to keep in mind that the scans were also evaluated during scanning. If a scan was deemed to be compromised, the scan was repeated if possible. In addition, participants who experienced problems lying in the scanner, and were more likely to move, were less likely to complete more than the first (T1w) scan, thus not being eligible for the study due to lack of the second (T2w) scan.

Still, to further assess potential confounding effects of image quality due to motion, we repeated our main analyses in two secondary analyses after excluding the 49 participants that scored below the 10^th^ percentile on two measures of image quality putatively related to head motion: (i) the temporal signal-to-noise ratio in diffusion weighted imaging data acquired in the same scanning session (Roalf et al. 2016); and (ii) and the Euler number (Rosen et al. 2018) estimated in the T1w images (see ***Supplementary Material*** for details).

T1w/T2w maps were created using the Human Connectome Project (HCP) processing pipeline (https://github.com/Washington-University/Pipelines), including processing with the Freesurfer 5.3 suite (http://surfer.nmr.mgh.harvard.edu). The T1w volume was divided on the aligned (using *bbregister* (Greve and Fischl 2009) and spline interpolation (Glasser and Van Essen 2011)) T2w volume, creating a T1w/T2w ratio volume. To estimate regional T1w/T2w, we used a multi-modal parcellation to divided each cerebral hemisphere into 180 areas (Glasser *et al.* 2016). We sampled T1w/T2w values from the WM/GM boundary, at 9 intermediate cortical depths (10% intervals), and at the GM/CSF boundary (***Supplementary Material, Fig. S2***). Here we focused on T1w/T2w measures at 70% depth from the pial surface to avoid partial volume effects (Huntenburg et al. 2017) (***Supplementary Material, Fig. S3***).

### Estimation of Growth Curves

To fit age trajectories without assuming a specific shape of the relationship *a priori*, we used penalized splines (Wood 2006; Fjell et al. 2010). Eight piecewise cubic B-spline basis functions were used to fit a smooth, non-linear curve from the weighted sum of these functions (***Supplementary Material, Fig.* S4**). We calculated the derivative, i.e. the slope at each point along the curve, as the differences (∆*y*) between the predicted curve (using the *predict* function in R (www.r-project.org/)) from i) one set of age values, and ii) another set with slightly increased age values, divided by this increment 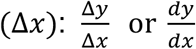. To find significant increases or decreases, we estimated the 99% CI of the derivative. The CIs were estimated using predict.gam (Marra and Wood 2012) in R (https://cran.r-project.org), with the Bayesian posterior covariance matrix (see Supplementary Material for further details). Based on the CI, we extracted 3 curve features, or milestones, namely age at peak growth, age at onset of stability, and age at onset of decline, for each region. We also estimated cross-sectional measures of the rate of growth, and the rate of decline (***Supplementary Material, Fig. S5***). As the CIs of the derivative curve are critical for the milestone estimates, we also calculated the CI using a bootstrap method to evaluate consistency across methods. Specifically, we drew 2000 random samples with replacements across participants, and calculated the growth curve and its derivative as above for each region. The 99% CIs were taken as the 0.05 and 0.995 percentile, respectively, of the resulting distribution of samples, at each point along the curve and the derivative, respectively. The percentile method was chosen for its simplicity compared with the original approach, although more complex, but potentially more accurate, alternatives have been proposed (DiCiccio and Efron 1996).

### Comparison with the Cytoarchitectonic Map of von Economo and Koskinas

We explored the relationship between cortical histology and the key milestones (Vertes et al. 2016). Based on previous work (Solari and Stoner 2011; Scholtens *et al.* 2016), we assigned each of the 360 regions to 1 of 5 cytoarchitectonic types classified according to the scheme of von Economo and Koskinas (Triarhou 2007), reflecting the 5 structural types of isocortex, or to two additional subtypes (Vertes *et al.* 2016): limbic cortex, and the insular cortex.

### Network Analyses

#### Structural network

To investigate how the milestones, which are local in nature, might relate to global network properties of the brain, we constructed an anatomical network using structural covariance analysis (Alexander-Bloch et al. 2013; Evans 2013). This approach allowed us to make a direct link, by using the same structural MRI data, between the milestone estimation and the network analysis. It has been shown that regions with high structural covariance or correlation are often involved in the same cognitive function and connected via white matter pathways (Lerch et al. 2006). Specifically, we correlated T1w/T2w for each region with all other regions, across participants between the milestones of maturity onset (34 years) and decline onset (72 years) in the global curve (see Fig. 1B), i.e., during the mature period with relatively stable T1w/T2w levels. This procedure yielded a 360×360 connectivity matrix, which was binarized employing a minimum spanning tree approach followed by global thresholding, retaining 10% of the strongest connections or edges (Supplementary Material, Fig. S6A (Alexander-Bloch et al. 2010). From this model, we assessed degree and modularity, two of the most common and, for degree, interpretable network metrics. These analyses were carried out in Matlab (https://www.mathworks.com) using the Brain Connectivity Toolbox (Rubinov and Sporns 2010). The community structure was obtained using the Louvain algorithm (Blondel et al. 2008), and consensus clustering (Sporns and Betzel 2016). We empirically defined the resolution parameter by finding a local minimum for nodal versatility of modular affiliation (Shinn et al. 2017) (see ***Supplementary material*** for details). A network representation of the structural connectivity (**Fig. 4ii**) was visualised using NetworkX, version 2.1 (https://networkx.github.io).

**Fig. 1.**
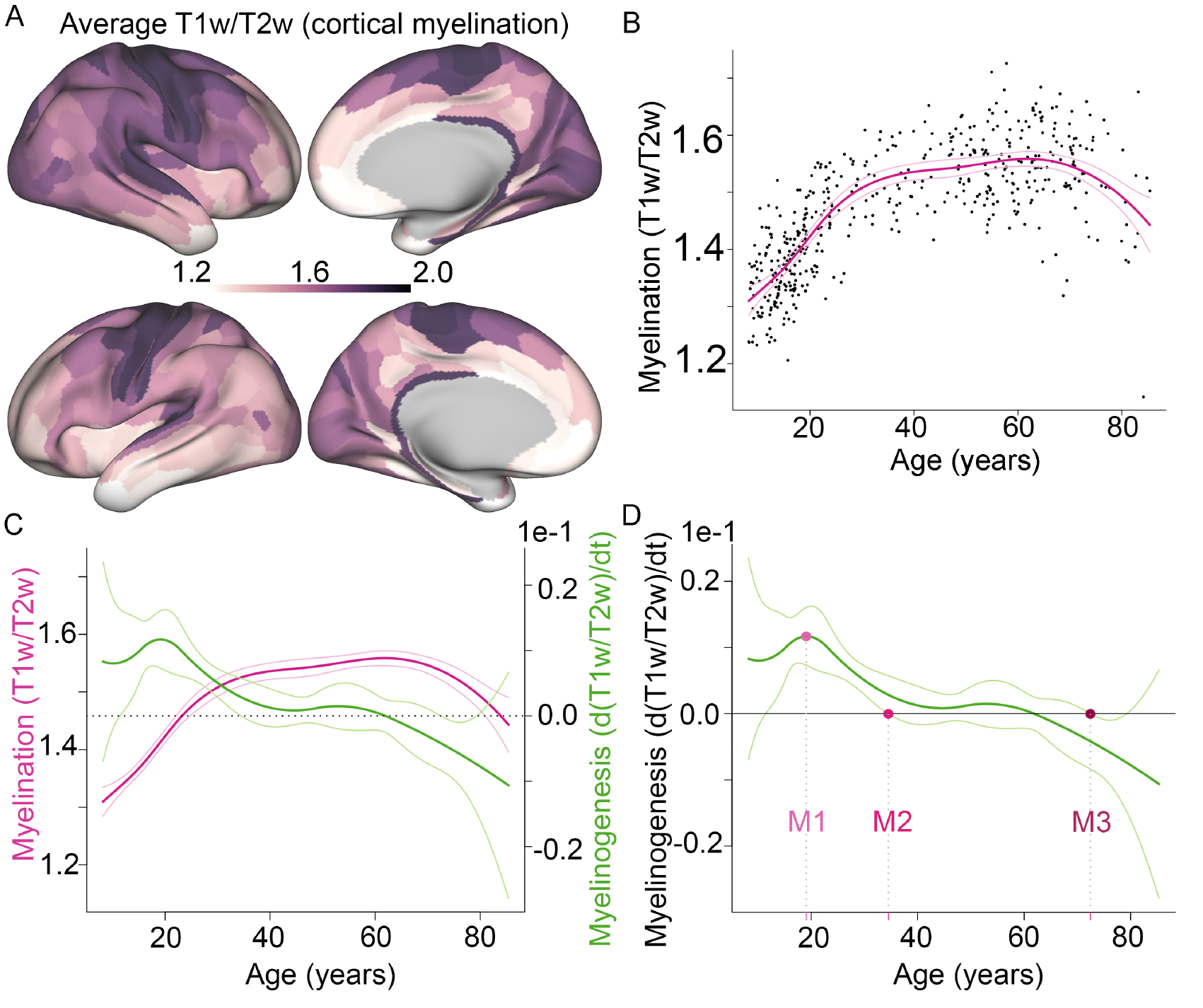
Milestones of human intra-cortical myelinogenesis estimated from non-linear growth curves of T1w/T2w MRI data measured across the life-cycle. **A**. Surface maps of T1w/T2w values averaged across the nearly 80 years of the lifespan (N=484). **B**. Average T1w/T2w values across the 360 regions for each participant plotted against the participants’ age. The fitted cubic spline and the 99% CI are plotted in magenta. **C.** The fitted cubic spline and the 99% CI (magenta) from **B**, and the estimated rate of change (first derivative) and the 99% CI (green). **D.** The 3 milestones: M1 = peak growth age (19 years), M2 = onset stability age (37 years), M3 = onset decline age (69 years).

#### Functional network

To further assess the relationship between the milestones, and brain network topology, we also created a functional network based on functional MRI scans at rest for a subset of the participants (aged between 20 and 72 years). The aim was to provide complementary information as structural and functional networks are robustly, but complexly related (see for instance Vincent et al. (2007)). Details of the sub-sample and the pre-processing can be found in the Supplementary material. From the resulting pre-processed fMRI volumes, regional average time series were extracted for the 360 regions. The time series were then fed to a wavelet decomposition (Bullmore et al. 2004) to 4 wavelet scales using the maximal overlap discrete wavelet transform with a Daubechies 4 wavelet, performed by use of the brainwaver package (Achard et al. 2012) in R (www.r-project.org/). We chose wavelet scale 1 frequency band (0.08-0.17 Hz) for further investigation, but also repeated the analyses at the scale 2 frequency band (0.04 −0.08 Hz) to assess consistency. The wavelet correlation between every pair of regions yielded one 360 × 360 matrix of regional functional connectivity per participant. We then averaged these matrices across participants resulting in one 360 × 360 matrix of regional functional connectivity. Similar network construction and analyses were performed as for the structural network, with focus on the total degree analysis. Thus, we constructed a binary graph by retaining the 10% strongest connections for this average functional connectivity network (***Supplementary Material, Fig. S7A***), using the minimum spanning tree approach. This network representation was fed to network analyses of community structure, and total degree centrality.

### Statistical Analyses

We tested for unimodality using Hartigans’ dip test statistic (Hartigan and Hartigan 1985). We used an expectation–maximization (EM) algorithm to fit Gaussian finite mixture models with 1 and 2 components, where a maximum of 2 components would provide proof of concept, and not be liable to over-fitting, and tested for best fit using a bootstrap likelihood ratio test with 10000 bootstraps. To facilitate the fitting process, 9 values above 25 years of age, and 4 values around 40 years were removed from M1 and M3, respectively, before estimation and plotting. Relationships between the regional measures were tested using Spearman’s rank correlation, except between rate of maturation and decline, where a linear regression was performed. To assess differences in rates of maturation and decline, respectively, between early and late waves across the 3 milestones, we calculated 99% CIs. In the cases where the CI overlapped, we tested for differences between waves by Wilcoxon rank sum test. The same approach was used to test for differences in the degree measures between waves. To test for the differences across cytoarchitectonic classes, we employed Kruskal-Wallis tests by rank, and accounted for the multiple tests performed by applying false discovery rate (FDR) correction (Benjamini and Yekutieli 2001).

## Results

### Trajectory and Milestones of Age-Related Differences in Micro-Structural MRI

The average T1w/T2w ratio map across all participants shows that highly myelinated primary motor and sensory areas have higher T1w/T2w values compared with association areas (**Fig. 1A**). The global trajectory of T1w/T2w, on average over all cortical regions in each participant, is approximately an inverted-U function of age over nearly 80 years of the lifespan (**Fig. 1B**). There is a maturational increase or growth of T1w/T2w in the first part of life, before the curve gradually levels off and a plateau is reached in middle age, followed by a later period of decline from around 60-65 years.

To quantify this age-related micro-structural variation more precisely, we estimated the slope of the T1w/T2w curve at each point in time, t, i.e., the first derivative of the trajectory, d(T1w/T2w)/dt (**Fig. 1C**). We then estimated 99% confidence intervals (CIs) for both the curve and its first derivative, which we used to rigorously define three key milestones (M1, M2, M3). M1, peak growth age, was defined as the age of maximum positive rate of change in T1w/T2w; M2, onset stability age, was defined as the earliest age after M1 at which the 99% interval for the derivative curve included zero, indicating that there was no significant growth or decline in T1w/T2w; and M3, onset decline age, was defined as the earliest age after M2 at which the 99% interval for the derivative curve did not include zero, indicating that there was significant negative rate of change in T1w/T2w (**Fig. 1D**). For the whole brain trajectory, peak growth age (M1) occurred at 19 years, onset stability age (M2) at 34 years, and onset decline age (M3) at 72 years. From the derivative curves we also estimated two additional metrics of interest (***Supplementary Material, Fig. S5***): the maximum rate of growth in T1w/T2w (i.e. the derivative at M1), and the maximum rate of decline (i.e. the derivative at mid-point between M3 and the end of the age range).

### Cortical Mapping of Milestones of Age-Related Differences in MRI Micro-Structure

We estimated all three milestones for each of 360 cortical regions and plotted the age at each milestone for each region as cortical surface maps. Peak growth age (**Fig. 2Ai**, and ***Supplementary Material, Fig. S8***) occurred earlier, i.e., before 13 years of age, predominantly in areas with high myelin content such as the primary motor and sensory areas, as well as the retrosplenial cortex and right LO2 (please see Glasser *et al.* (2016)for explanation of the nomenclature), and area MT. Regions with a later peak growth age, i.e., after 13 years of age, were found in prefrontal, parietal, and temporal cortices, particularly in the right hemisphere, as well as medial occipital cortex.

**Fig. 2.**
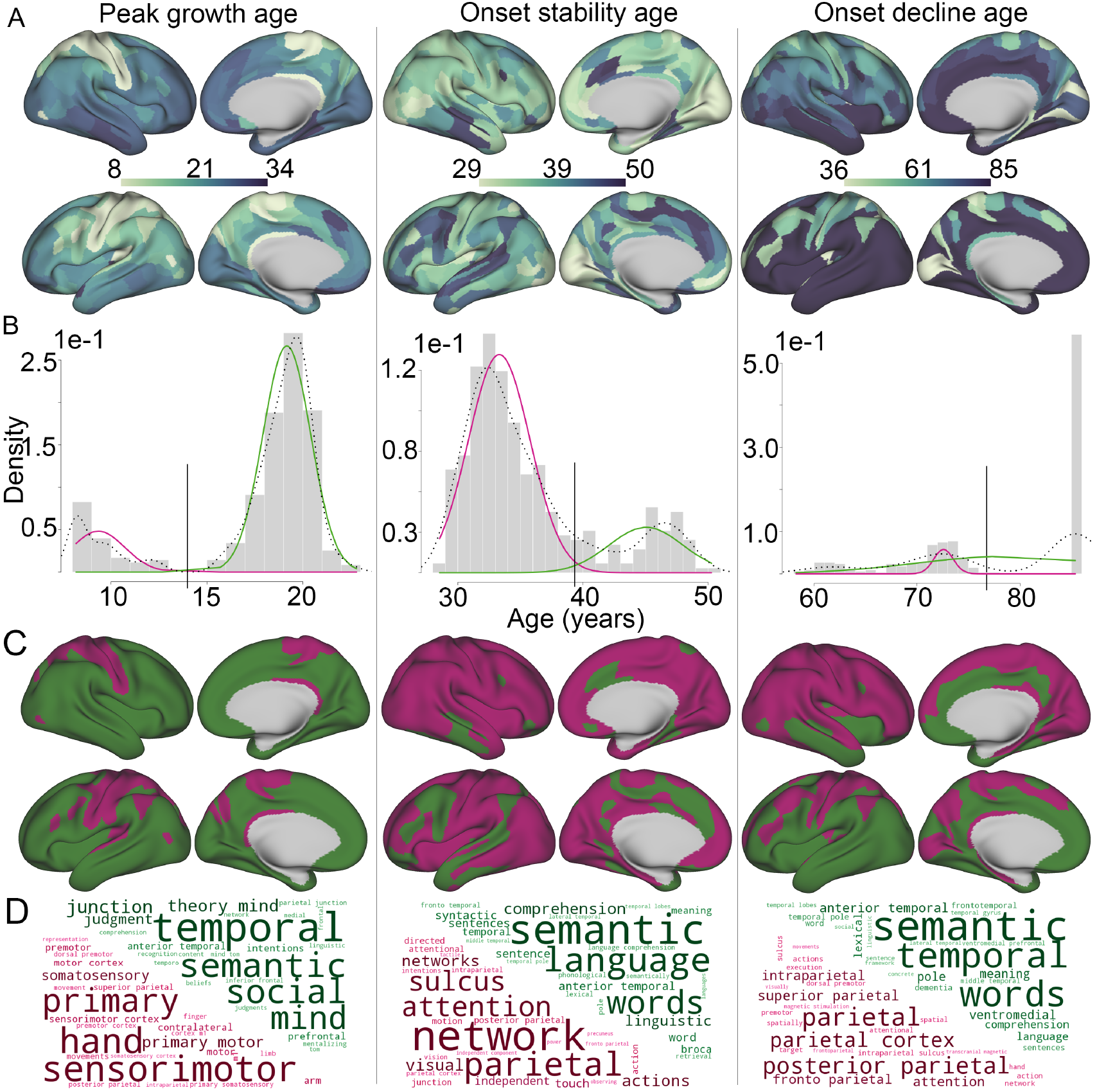
Intracortical T1w/T2w MRI milestone maps, and bimodal distributions suggesting that timing of generation and degeneration of myelination occurs in waves. **A**. Surface maps of peak growth age, onset stability age, and onset decline age. **B**. Histograms for i) peak growth age, ii) onset stability age, and iii) onset decline age with the fit of a 2-component Gaussian mixture model overlaid (in magenta and green). Age used for dichotomisation into early and late waves (grey vertical bar) for peak growth age was 12.7 years, stability 39.5 years, and decline 77 years. Black dotted line = probability density. **C**. Surface maps of early (magenta) and late (dark green) waves based on **B**. **D**. Word clouds based on correlations with NeuroSynth meta-analysis maps for the early and late waves at each milestone.

Onset of stability was reached between 29 and 50 years (**Fig. 2Aii**) and was found to occur later particularly in lateral temporal, medial prefrontal and anterior cingulate cortex bilaterally, as well as in left lateral frontal cortex, with regions in general reaching a plateau later in the left hemisphere.

Onset of significant decline started between 58 and 77 years of age (**Fig. 2Aiii**) in 170 regions, while 190 regions, particularly in the left hemisphere, did not show a significant decline. In both hemispheres, onset of decline was observed earlier in motor and sensory cortices, and the retrosplenial complex.

### Bimodal Distributions of Micro-Structural Milestones

Inspection of the distributions of each of the milestones provides intuitive support for the existence of waves of change in T1w/T2w. For each milestone, there are two distinct peaks (**Fig. 2B**), suggesting that the distribution is not unimodal but at least bimodal (see also quantile-quantile plots, ***Supplementary Material, Fig. S9*)**. We tested this observation formally, finding evidence for significant non-unimodality in all milestone distributions (Hartigan’s dip test, *P* < 0.001). A two-component mixture of Gaussian distributions was fitted to each of the milestone distributions, providing a significantly better fit than a unimodal model in all cases (bootstrap likelihood ratio test, all *P* < 0.001). Each milestone distribution could thus be partitioned into an early wave and a late wave (**Fig. 2B)**, and each cortical region could be assigned to the early or late waves of each milestone (**Fig. 2C**).

The first wave of peak growth occurred at a mean age of 9.4 years ([95% CI [9.0, 9.7]) in relatively few areas of primary cortex, specialised for motor and somatosensory function; the second wave occurred at a mean age of 19.5 years (95% CI [19.2-19.7]) in a larger number of association areas, specialised for semantic and social functions, such as theory of mind (determined through a search of neurosynth.org (Yarkoni et al. 2011), **Fig. 2D**).

The first wave of onset of stability occurred at a mean age of 33.3 years (95% CI [33.0, 33.6]), and involved areas of parietal cortex specialised for attention. A smaller number of areas, mostly located in the left hemisphere and specialised for semantic language functions, reached stability later with mean age of onset 45.1 years (95% CI [44.4, 45.8]).

The first wave of onset of decline occurred at a mean age of 68.8 years (95% CI [67.8, 69.8]), and involved mainly right hemisphere frontal and parietal regions specialised for attention and executive functions. The second wave in this case represented a large number of regions, mainly in the left hemisphere and specialised for semantic language functions, which did not show evidence of significant decline within the age range of the sample (up to 85.4 years).

### Rates of Maturation and Decline of Cortical Micro-Structure

Surface maps of the regional variation in rate of maturation, and the rate of decline, are shown in **Fig. 3A** and ***Supplementary Material, Fig. S10***. We observed higher rates of maturation and decline in highly myelinated areas of primary motor and sensory areas and lower rates of maturation and decline in lightly myelinated frontal and temporal cortical areas. The average T1w/T2w map (**Fig. 1A**) and the rate of peak growth map (**Fig. 3Ai**) were strongly positively correlated (Spearman’s ρ = 0.75, *P* < 0.001). Conversely, the average T1w/T2w map was strongly negatively correlated with the rate of decline map (**Fig. 3Aii**; Spearman’s ρ = −0.67, *P* < 0.001). As expected given these results, the regions with high rates of maturation also had high rates of decline (**Fig. 3B**; Spearman’s ρ = −0.59, *P* < 0.0001), i.e. a “fastest in, fastest out” pattern.

**Fig. 3.**
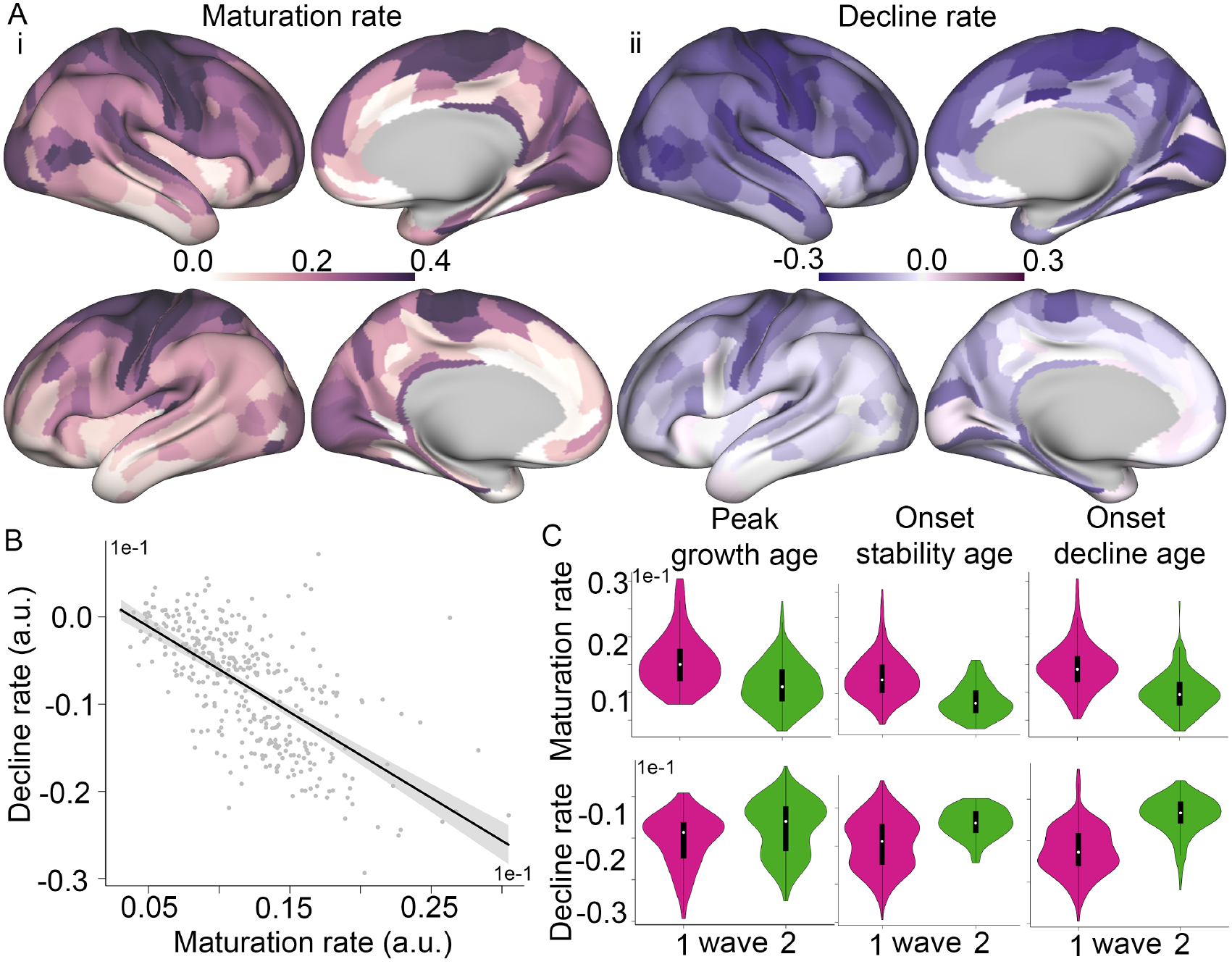
Milestones of intracortical MRI growth curves were related to cross-sectional rates of change in generation and degeneration of myelination. **A**. Surface maps depicting rates of i) peak growth (i.e. the derivative at M1), and ii) decline (i.e. the derivative at mid-point between M3 and the end of the age range). **B**. Inter-regional association between micro-structural maturation, and decline rate. Regression line in black (R^2^ = 0.43, p < 0.001), the 95% CI in shaded grey. **C**. Violin plots of the relationship between rate of peak growth (top row), and decline (bottom row) with age of peak maturation (first column), stability, decline, respectively, showing significant differences between early (magenta) and later (green) waves at each milestone.

These maturation and decay rates of T1w/T2w, respectively indicative of rates of cortical myelination and demyelination, were significantly different between waves as defined by the three milestones, M1-M3 (**Fig. 3C**). The rate of maturation was significantly more positive, and the rate of decline was significantly more negative, in the first M1 wave of regions that reached peak growth before puberty than in the second M1 wave of regions that reached peak growth after puberty (***Supplementary Material, Table S1A***). Likewise, the first M2 wave of regions, which reached onset of stability earlier, had more positive rates of maturation than the second M2 wave of regions, which reached stability later. Finally, a similar pattern was evident for the M3 milestone: regions in the first wave, which reached onset of decline earlier, had more positive rates of maturation. Regions in the second M3 wave had less positive rates of maturation and less negative rates of decline (close to zero).

### Onset decline age and cortical thickness

The findings presented above for onset stability age (M2) and onset decline age (M3) point to a “last in, last out” pattern. That is, in general the regions which reach stability later (in the second wave of M2) are more likely to also show later onset of age-related decline (in the second wave of M3). This result is partly at odds with the “last in, first out” hypothesis, which is supported by studies using macro-structural measures, such as cortical thickness. Thus, we tested the relationships between age and cortical thickness in the early and late waves of onset decline age. To this end, similarly to previous analysis for the “last in, first out” hypothesis (Raz 1999), we calculated the regression slope F value (weighted by direction of the slope as 11 slopes were not negative) of the cortical thickness-age relationships in each region, and interpreted more negative age-relationships as indicative of “first out”. That is, the “first out” manifests as steeper age-related decline.

The sample used was formed of participants aged 34.5 (onset stability age for the average curve, Figure 1C) or older (n=242). Then, we tested whether the weighted F values were different for the two waves of onset of decline age. As hypothesized based on Raz (1999), regions in the second wave showed a stronger decline with age compared to regions which reached maturity earlier (Wilcoxon rank sum test W = 21194, P < 0.0001). This relationship was not altered by taking the T1w/T2w level of the region into account. This pattern is accordance with the “last in, first out” hypothesis. That is, the regions reaching onset of stability age late (“last in”), showed stronger decline in cortical thickness with higher age (“first out”).

### Cytoarchitectonics

We tested the relationship between cortical histology and milestones of the T1w/T2w trajectory by classifying each of the 360 regions according to its membership of one of 7 cytoarchitectonic classes defined by the von Economo and Koskinas atlas (Scholtens *et al.* 2016) (***Supplementary Material, Fig. S11A***). Different cytoarchitectonic classes and cerebral hemispheres were differently represented in the two waves at the M1 milestone (***Supplementary Material, Fig. S11*B**). The proportion of regions reaching peak growth before puberty was significantly different between cytoarchitectonic classes, with a higher proportion of primary motor and primary sensory cortices in the first wave than in the second wave (χ^2^= 65, df=6, *P*_FDR_ = 1× 10^-10^). The differences in proportion of cytoarchitectonic classes in the first and second waves of the M2 milestone were not significant after multiple comparison correction (χ^2^= 14, df=6, *P*_FDR_ = 0.126, *P*_uncorrected_ = 0.035). However, there were significant differences between the first and second waves of the M3 milestone, with a higher proportion of association cortical areas, especially in the left hemisphere, represented in the second wave (χ^2^= 37, df=6, *P*_FDR_ = 1.3 × 10^-5^).

### Connectomics

To investigate how the local cortical milestones of age-related differences in T1w/T2w might relate to the topology of the cortical connectome, we estimated the pair-wise correlation of T1w/T2w between each possible pair of regions across participants between the ages of stability and decline onset in the global curve (see **Fig. 1B**; 37 and 69 years, respectively), that is, a period with relative stable T1w/T2w levels. From this micro-structural network model (***Supplementary Material, Fig. S6A***), we constructed a binary graph by retaining the 10% strongest connections which could be decomposed into a community structure of modules (***Supplementary Material, Fig. S6B***). For each regional node, we then estimated its degree centrality (Bullmore and Sporns 2009), i.e., the number of edges connecting it to the rest of the network; its intra-modular degree, i.e., the number of edges connecting it with nodes in the same module; and its inter-modular degree, i.e., the number of edges connecting it with nodes in other modules. The network had a fat-tailed degree distribution (***Supplementary Material*, Fig. S6C**), indicative of so-called hub regions. These high degree regional nodes were mainly found in frontal, parietal, and temporal association cortices (**Fig. 4Ai-ii**).

**Figure 4.**
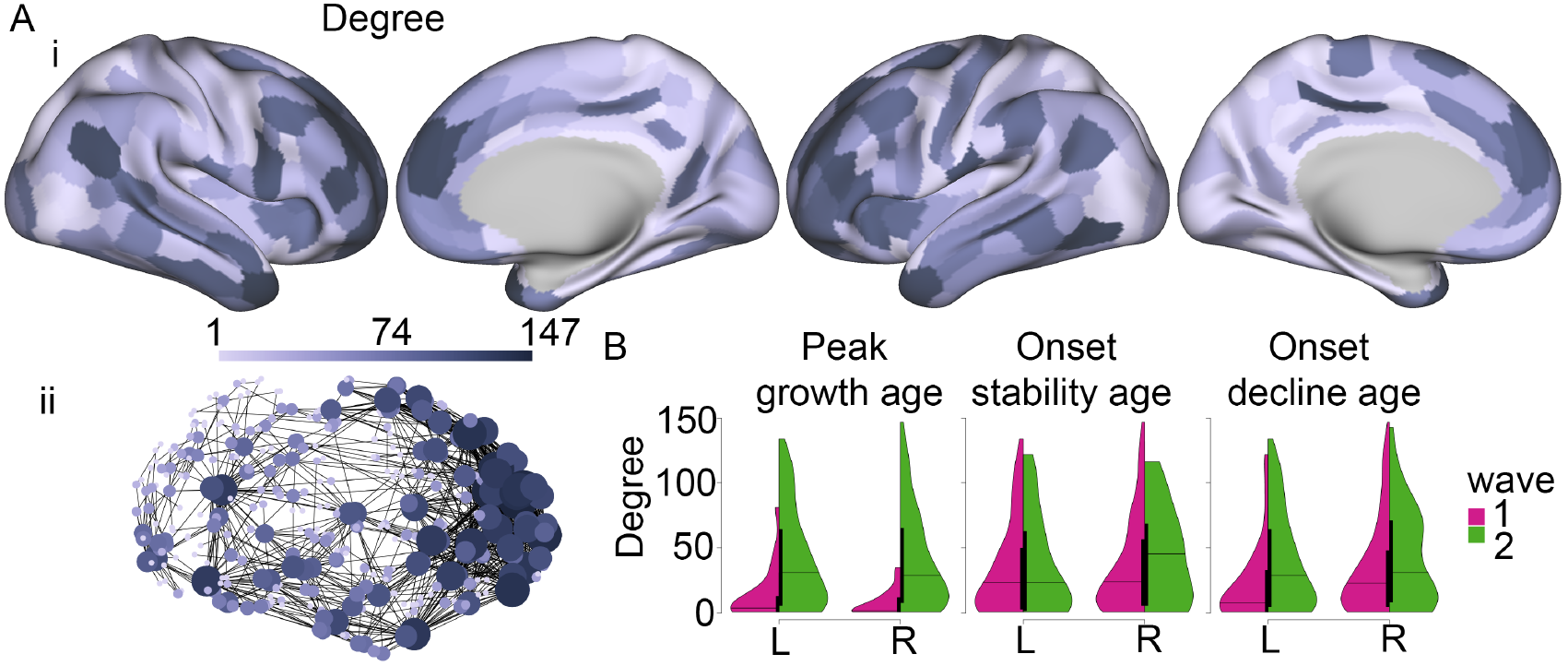
Hubs of the structural covariance connectome have a long period of adolescent and early adult myelination, with delayed onset of maturity, but later onset of decline. **A**. The structural covariance network. High-degree centrality hubs were concentrated anatomically in association cortical areas. i. Surface maps of degree. ii. Network representation, nodes are coloured and sized by degree (only the top 2% strongest connections are shown for clarity). **B**. Violin plots, split per hemisphere and early (magenta) and late (green) waves, for peak growth age, onset stability age, and onset decline age. L/R = left/right hemisphere.

There were significant differences in nodal topology between the two waves of the M1 milestone (**Fig. 4B**, and ***Supplementary Material, Fig. S12*** and ***Table S1B***). Regions which reached peak growth before puberty had significantly reduced total degree, intra-modular degree, and inter-modular degree, compared to regions which reached peak growth after puberty. Likewise, there were significant differences in nodal topology between the two waves of the M3 milestone. Regions in the second wave, which did not show onset of significant decline, had significantly higher total degree, and inter-modular degree, than regions in the first wave which had onset of decline in the 60s. There were no significant differences in nodal topology between regions in the first and second waves of the M2 milestone.

The functional network could also be decomposed into a community structure of modules (***Supplementary Material, Fig. S7B***), and had a fat-tailed degree distribution (***Supplementary Material, Fig. S7C***). The high degree regional nodes were mainly found in primary visual and motor cortices, as previously reported (Crossley et al. 2013) (***Fig. 5Ai-ii***).

**Figure 5.**
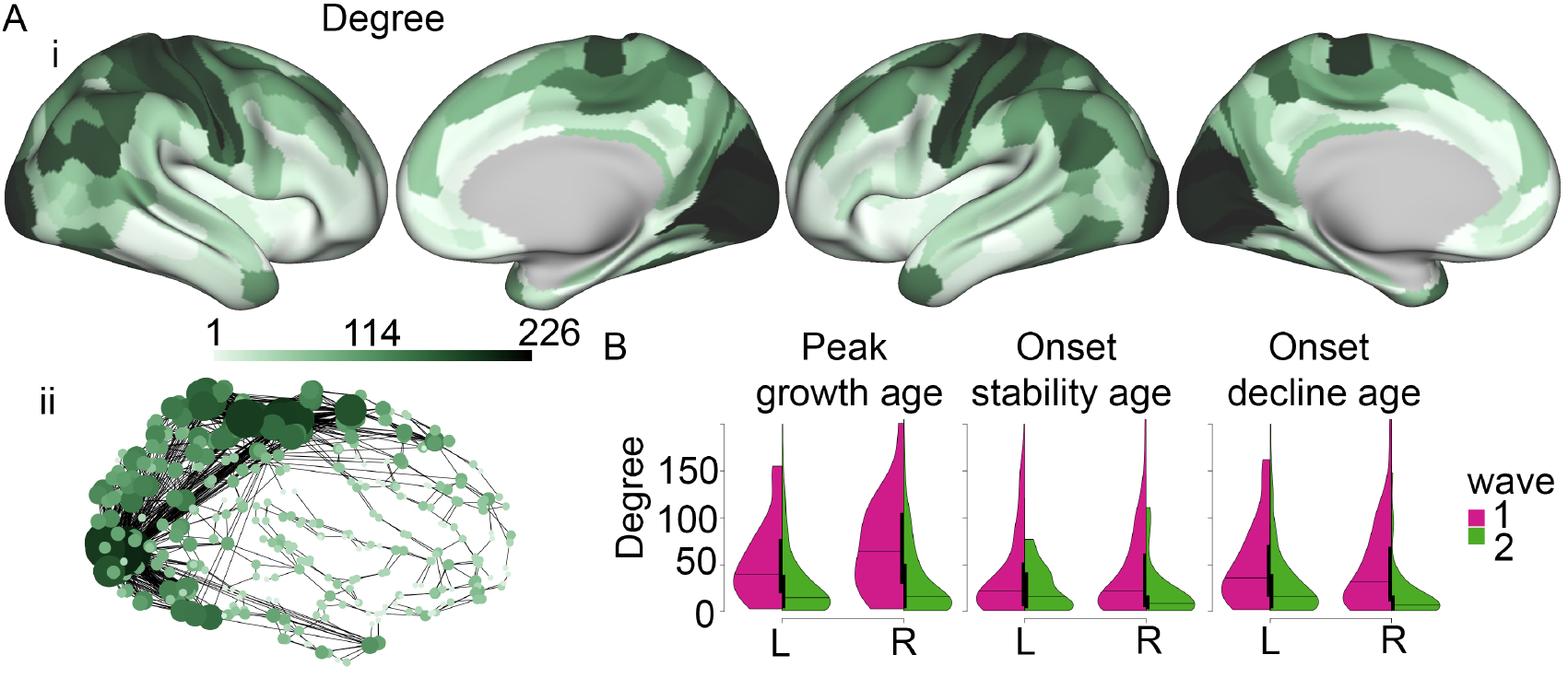
Hubs of the functional connectome have a short period of adolescent and early adult myelination, with early onset of maturity, and early onset of decline. **A.** The functional network. High-degree centrality hubs were mainly concentrated in primary visual and motor cortices. i. Surface maps of degree. ii. Network representation, nodes are coloured and sized by degree (only the top 2% strongest connections are shown for clarity). **B.** Violin plots, split per hemisphere and early (magenta) and late (green) waves, for peak growth age, onset stability age, and onset decline age. L/R = left/right hemisphere.

As for the structural network, there were significant differences in functional network nodal topology between the two waves of the milestones (***Fig. 5B***, and ***Supplementary Material, Table S1B***). However, as functional network hubs were mainly found in primary visual and motor cortices, the patterns were different, with early wave regions showing higher total degree. For M1, regions which reached peak growth before puberty had significantly higher total degree compared to regions which reached peak growth after puberty. Likewise, there were significant differences in nodal topology between the two waves of the M3 milestone: regions which had onset of decline in the 60s had significantly higher total degree than regions which did not show significant age-related decline. In contrast to the structural network, there was also a significant difference in nodal topology between regions in the first and second waves of the M2 milestone. Again, regions in the first wave, which reached stability earlier, had significantly higher total degree than regions in the second wave which reached onset of stability later. Repeating the analysis using the wavelet scale 2 frequency band (0.04 −0.08 Hz) produced nearly identical results (***Supplementary Material, Table S1B***).

#### Sensitivity analyses

We repeated parts of our analysis by calculating the CIs based on the bootstrap method. As the CIs based on the bootstrap method were highly similar to the original CIs (***Supplementary Material, Fig. S14Ai-ii***), the milestones based on the bootstrapped CIs were also highly similar (***Supplementary Material, Fig. S14Aiii-v***).

We also performed two additional analyses based on sub-samples of 435 participants with 49 participants excluded on the grounds that they were in the lowest 10% of the sample for image quality metrics. For the analysis where 49 participants were excluded based on the tSNR, the fitted spline curve was highly similar to the curve estimated on the full sample (Pearson’s r = 0.98, ***Supplementary Material, Fig. S14Bi-ii***). For the analysis where participants were excluded based on the Euler number, the fitted curve in the younger half of the sample was again very similar to the full sample (r = 0.95, ***Supplementary Material, Fig. S14Ci-ii***). For the latter part of the age range, there was less evidence of a decline in T1w/T2w ratio (for 24 regions compared to 190 regions) compared to the results of analysis on the full sample. This reduction in regions showing decline when data were quality controlled by the Euler number criterion also made the distribution for M3 less clearly non-unimodal than it was in the whole sample (Hartigan’s dip test, P = 0.02 versus P < 0.001, ***Supplementary Material, Fig. S14Cv***). For the other milestones, the histograms of both quality-controlled sub-samples showed a non-unimodal distribution similar to the distributions identified by analysis of the whole sample (***Supplementary Material, Fig. S14Biii-v***, and ***S14Ciii-iv).***

## Discussion

Based on a cross-sectional sample covering nearly 8 decades of the lifespan, we delineated the age-related trajectory of maturation and senescence of a micro-structural MRI marker linked to myelin. We found indications of maturation and decay occurring in stages, or waves, at three key milestones: age at peak growth, age at onset of stability, and age at onset of decline. Importantly, we also observed that regions in the first wave of developmental milestones had faster rates of maturation and faster rates of decline. This result contradicted the “last in, first out” hypothesis (Raz 1999), according to which early maturing regions should be more protected from age-related decline than late maturing regions. We found the opposite. By linking the timing of milestones to the cytoarchitectonic atlas of von Economo and Koskinas, we found that early maturing regions were histologically classified mainly as heavily myelinated primary motor and sensory cortices, whereas later maturing regions were mainly association, limbic and insular cortices. We also found that the high degree hubs of the anatomical connectome had significantly later age at peak growth and later age at onset of decline than less well-connected nodes in the structural covariance network.

### Waves of Myelinogenesis as Implicated by Age Differences in Milestones

We observed a bimodal distribution of all milestones. In accordance with histological results based mostly on early childhood brains (Flechsig 1901; Yakovlev and Lecours 1967), we interpret this finding as suggestive of waves of myelinogenesis. This concept has not been studied within a lifespan perspective, and the observations could be conceptualized as “echoes” in later life of earlier waves of neuro- and glio-genesis (Rakic 2002). Another interesting hypothesis relates post-pubertal myelination to sexual maturity (Miller et al. 2012), and the changes in executive and social cognition associated with adolescence (Blakemore and Choudhury 2006). This suggestion relates to the timing of peak myelination which occurred in one wave before and one wave after puberty. The post-pubertal wave regions were more concentrated in association, insular, and limbic cortices, and this accords with the suggested link between post-pubertal myelination and the onset of psychiatric diseases during adolescence and early adulthood (Bartzokis 2002; Paus et al. 2008).

### The “Last In, First Out” Hypothesis

The present micro-structural results indicated a “last in, last out” pattern. This result is in line with a recent study in marmosets using a similar approach to the one taken here, reporting that GM volume in primary motor and sensory regions generally declined earlier than associations regions (Sawiak et al. 2018). In contrast, the macro-structural cortical thickness results were in line with the “last in, first out” hypothesis (Raz 1999). MRI lifespan studies using grey matter (GM) macro-structural measures have found support for this hypothesis (e.g. Douaud et al. (2014)), while others using micro-structural measures of the cortex (e.g. Westlye et al. (2010)) have not. One possible explanation for this discrepancy between the lifespan trajectories for macro-versus micro-structural aspects of the cortex may be that loss of myelin in primary motor and sensory areas could offset cortical thinning. As both the cortex and the subjacent white matter become “greyer” due to demyelination, the WM/GM boundary recedes, making thinning appear less extensive (Westlye et al. 2009). This process is essentially an inversion of the suggested explanation of thinning in childhood and adolescence, where increased myelination makes the deeper layers of cortex appear “whiter” (Seldon 2005). The more negative macro-structural relationships with age observed in the lightly myelinated late areas might be driven more by processes other than demyelination, for instance reduction in dendritic spine density of pyramidal neurons (Kabaso et al. 2009). Another possible explanation is that demyelination with age is offset by other types of myelin degeneration, such as the formation of redundant myelin and increasing thickness of myelin sheaths, as reported in older aged monkeys (Peters 2009).

### The Gain-Predicts-Loss Hypothesis

Although we did not find a clear relationship between age at onset of stability and age at onset of decline, we did find that regions showing higher rates of peak growth, and doing so relatively early, also showed an early onset of decline and a faster rate of decline. The mainly primary motor and sensory regions showed earlier and faster rates of cortical myelination and earlier and faster rates of cortical demyelination with high age. Thus, in addition to a “last in, last out” pattern, our results fit partly with a “gain-predicts-loss” hypothesis (Yeatman *et al.* 2014). This hypothesis predicts that the rate of change in development will be mirrored by the rate of decline in aging. We found that the aging process is not exactly the reverse of development, in contrast to R1 trajectories of WM tracts (Yeatman *et al.* 2014). Instead, the T1w/T2w trajectories increased more steeply in development compared with less rapid rates of decline later in life, as in the diffusivity trajectories of the same tracts (Yeatman *et al.* 2014). Thus, there is a similarity between the models of micro-structural development over the lifespan in WM and cortex. Whether these effects are governed by similar or different mechanisms warrants closer scrutiny in future studies. Interestingly, the differences in rate of development and decline in primary motor and sensory regions compared with association, limbic and insular regions, fits well with the processing-speed theory of adult age differences in cognition (Salthouse 1996). This theory proposes that processing speed reduction leads to the age-related decline in higher-order cognition, such as memory and reasoning. In development, a similar idea, but in an opposite direction with increased speed, has been proposed (Kail 1991). Accordingly in this context, processing speed relates more strongly to sensorimotor areas than association cortices, while association cortices relate more strongly to higher-order cognitive functioning (Deary et al. 2010).

### Connectomics

Another feature of our findings was the relationship between waves of putative (de-)myelination and high degree network regions of the anatomical connectome, so-called hubs. We found that structural hubs had significantly later age at peak growth and later age at onset of decline than less well-connected nodes in the structural covariance network. Flechsig (1901)observed that association regions were the focus of a relatively late wave of myelinogenesis. In our data and previous studies (Alexander-Bloch *et al.* 2013; Whitaker *et al.* 2016), structural hub regions are disproportionately located in association cortex. Thus, the observation of different micro-structural trajectories for hubs compared with primary motor and sensory areas aligns with Flechsig’s idea. Here, we extend this notion by showing that structural hub regions seem to be the focus of a relatively late wave of myelinogenesis also later in life. Although the relationships were not strong, the result followed from our earlier findings that association cortices (and limbic and insular cortices) show later and lower rates of micro-structural maturation and decline compared with motor and sensory cortex. As discussed in Whitaker *et al.* (2016), cortical myelination in development might be topologically targeted to optimize structural hub performance. Similar optimization processes might be at play in aging, which might break down in disorders such as Alzheimer’s disease (Crossley et al. 2014).

We also found relationships between waves of putative (de-)myelination and the hubs of the functional network. However, the pattern here was different, as the functional hubs were more posteriorly located, particularly in primary visual and motor cortices, which mainly comprised the first wave of earlier maturing regions at each milestone. Compared to structural network hubs, the interpretation of degree-based hubs in functional networks hubs is less straightforward, as functional connections reflect indirect communication occurring within the underlying structural network (van den Heuvel and Sporns 2013). These findings should thus be interpreted with caution.

### Methodological Issues

First, the results are based on cross-sectional analyses, and longitudinal data are needed to estimate measures of individual change (Lindenberger et al. 2011). Cross-sectional studies may suffer from cohort effects, potentially exacerbated by differences in recruitment bias between age groups (Fjell et al. 2014). However, few longitudinal studies cover the lifespan, impeding measurements of change rates as a function of age, and such studies may also be influenced by selective attrition (Fjell *et al.* 2014). Longitudinal lifespan studies, measuring change over relatively few years (< 10 years) for each participant, may also suffer cohort effects. Hence, given the relative paucity of studies including most of the lifespan (i.e. from children to older adults), both cross-sectionally and longitudinally, we believe the current approach has some merits.

Second, the T1w/T2w ratio measure used here stems from T1- and T2-weighted scans, at a conventional resolution. Using a quantitative measure at high resolution (Huntenburg *et al.* 2017) likely would have been preferable for sensitivity and specificity. Moreover, the link between the T1w/T2w ratio and myelin content has recently been questioned. Indeed, it seems unlikely that the T1w/T2w ratio, or other MRI indices related to myelin such as R1 (Sigalovsky *et al.* 2006), should depend solely on myelin given the complex biophysical process underlying MRI relaxation rates. More specifically, Ritchie et al. (2018)reported positive correlations between the T1w/T2 ratio and myelin-associated genes, supporting its use as a myelin marker, but concluded that molecule size, transcription marking oligodendrocytes, axon calibre, and pH were more strongly associated with T1w/T2w. Arshad et al. (2017), in 20 participants aged between 24 and 70 years, observed positive correlations between T1w/T2w and myelin water fraction within 6 WM tracts, but negative correlations between different tracts. The authors suggest that the T1w/T2w ratio relates more to variation in calibre and packing density of the axons than myelin content. In this regard, it is interesting to note that R1, in both ex vivo and in vivo imaging of 6 and 4 WM regions, respectively, correlated with histology-derived axon size, but not with myelin content in rats (Harkins et al. 2016). Harkins *et al.* (2016)caution against interpreting T1 differences between regions or participants as necessarily reflecting differences in myelin content, even when using potentially more promising T1 measurements than used here to quantitatively map myelin content (Stuber et al. 2014). Accordingly, as previously noted (Westlye *et al.* 2010), interpretation of signal intensity differences across the lifespan should be made cautiously without histology data, as a host of neuronal and non-neuronal processes likely play out and potentially interact at various intensities and stages across development and aging (Tau and Peterson 2010; Lopez-Otin et al. 2013).

Interestingly, Righart et al. (2017)combined T1w/T2w imaging and histology, in 9 late stage secondary progressive multiple sclerosis patients aged 51 to 86 years. They correlated T1w/T2w ratio values in 5 normal-appearing cortical regions with histology-derived measures of myelin, axon and dendrites. The result showed that T1w/T2w ratio related significantly to dendrite, but not myelin (or axonal), density. Although, as acknowledged by the authors, the nature of the sample (late stage multiple sclerosis brain donors) might impede the generalisability to healthy participants, the finding is potentially important, and highlights the importance, and difficulty, of histological validation of in vivo MRI indices. Taken together, it might be more appropriate here, to use the term “apparent myelin”, similar to “apparent cortical thickness” and “apparent diffusion coefficient”, to highlight the aforementioned challenges in in vivo MRI (Walhovd et al. 2017).

Third, many regions (particularly in the left hemisphere) did not exhibit age-related decline, which contrasts with other MRI markers such as cortical thickness. Hemispheric asymmetry in patterns of age-related cortical volume shrinkage in aging have been inconsistent (Raz et al. 2004). Still, verbal abilities are relatively well preserved in aging (Shafto and Tyler 2014), and a potential relationship between intracortical micro-structure and preservation of verbal abilities, which are known to depend mainly on the left hemisphere (Herve et al. 2013), constitutes an interesting hypothesis for future research. Similarly, assessment of relationships between individual differences in the timing of these milestones and behavioural or neuropsychological changes associated with development would be an important task for future research.

Finally, recent studies have shown that even subtle motion might bias anatomical estimates in studies of brain structure and age (see for instance Alexander-Bloch et al. (2016)). Here, all images were thoroughly visually inspected for motion, both during and after scanning. Still, we further probed our data by excluding 10% of the sample with the lowest data quality defined by two criteria of image quality: tSNR from diffusion-weighted scans and the Euler number from the T1w scans. Repeating the main analyses on these sub-samples, the results in the younger half of the age range were virtually unaltered. This consistency of results was also apparent in the latter part of the age range when using tSNR as the quality index. However, when using the Euler number, the number of regions showing decline in older age was reduced to 24 regions compared with 170 regions in the full sample. However, it is unknown if this reduction in regions showing age-related decline in T1w/T2w ratio is due to head movement. For instance, it is possible that the Euler number could be increased in older adults with negligible motion during scanning because it is a measure of the topological complexity of the reconstructed cortical surface (Dale et al. 1999; Rosen et al. 2018) and age-related change in cortical structure could adversely impact on the performance of the cortical segmentation algorithm in FreeSurfer.

The consistency of results in the younger half of the sampled age range, where head motion has been identified as pervasive (see for instance Satterthwaite et al. (2012)), and in aging when using tSNR as quality index, supports the notion that our principal findings are not unduly influenced by movement during scanning. That said, these analyses were all post-hoc. Future lifespan studies of age-related structural brain effects should aim for direct and prospective estimates of head motion during scanning.

## Summary

As indicated by the present cross-sectional age differences, peak growth, stability, and decline of an MRI measure related to intracortical myelination proceeds in stages, or waves, in different regions. Primary motor and sensory regions reached all milestones earlier than association, limbic, and insular regions. The results contrasted with the so-called “last in, first out” hypothesis. Instead, the findings suggested a “last in, last out” pattern, where the regions less likely to show decline in aging were slowly maturing, association cortical hubs. In addition, the results fit partly with a gain-predicts-loss hypothesis, where regions which mature fastest, decline fastest. Future studies might benefit from looking more closely into how timing of waves relate to the onset of neuropsychiatric disorders, as well as the changes in cognitive and social skills, in development and aging.

## References

Achard S, Delon-Martin C, Vertes PE, Renard F, Schenck M, Schneider F, Heinrich C, Kremer S, Bullmore ET. 2012. Hubs of brain functional networks are radically reorganized in comatose patients. Proc Natl Acad Sci U S A. 109:20608–20613.

Alexander-Bloch A, Clasen L, Stockman M, Ronan L, Lalonde F, Giedd J, Raznahan A. 2016. Subtle in-scanner motion biases automated measurement of brain anatomy from in vivo MRI. Hum Brain Mapp. 37:2385–2397.

Alexander-Bloch A, Giedd JN, Bullmore E. 2013. Imaging structural co-variance between human brain regions. Nat Rev Neurosci. 14:322–336.

Alexander-Bloch AF, Gogtay N, Meunier D, Birn R, Clasen L, Lalonde F, Lenroot R, Giedd J, Bullmore ET. 2010. Disrupted modularity and local connectivity of brain functional networks in childhood-onset schizophrenia. Front Syst Neurosci. 4:147.

Arshad M, Stanley JA, Raz N. 2016. Adult age differences in subcortical myelin content are consistent with protracted myelination and unrelated to diffusion tensor imaging indices. Neuroimage. 143:26–39.

Arshad M, Stanley JA, Raz N. 2017. Test-retest reliability and concurrent validity of in vivo myelin content indices: Myelin water fraction and calibrated T1 w/T2 w image ratio. Hum Brain Mapp. 38:1780–1790.

Bartzokis G. 2002. Schizophrenia: breakdown in the well-regulated lifelong process of brain development and maturation. Neuropsychopharmacology. 27:672–683.

Bartzokis G, Beckson M, Lu PH, Nuechterlein KH, Edwards N, Mintz J. 2001. Age-related changes in frontal and temporal lobe volumes in men: a magnetic resonance imaging study. Archives of general psychiatry. 58:461–465.

Benjamini Y, Yekutieli D. 2001. The control of the false discovery rate in multiple testing under dependency. Annals of statistics. 1165–1188.

Blakemore SJ, Choudhury S. 2006. Development of the adolescent brain: implications for executive function and social cognition. J Child Psychol Psychiatry. 47:296–312.

Blondel VD, Guillaume JL, Lambiotte R, Lefebvre E. 2008. Fast unfolding of communities in large networks. J Stat Mech-Theory E.

Bock NA, Hashim E, Kocharyan A, Silva AC. 2011. Visualizing myeloarchitecture with magnetic resonance imaging in primates. Annals of the New York Academy of Sciences. 1225 Suppl 1:E171–181.

Bock NA, Kocharyan A, Liu JV, Silva AC. 2009. Visualizing the entire cortical myelination pattern in marmosets with magnetic resonance imaging. J Neurosci Methods. 185:15–22.

Bullmore E, Fadili J, Maxim V, Sendur L, Whitcher B, Suckling J, Brammer M, Breakspear M. 2004. Wavelets and functional magnetic resonance imaging of the human brain. Neuroimage. 23 Suppl 1:S234–249.

Bullmore E, Sporns O. 2009. Complex brain networks: graph theoretical analysis of structural and functional systems. Nat Rev Neurosci. 10:186–198.

Callaghan MF, Freund P, Draganski B, Anderson E, Cappelletti M, Chowdhury R, Diedrichsen J, Fitzgerald TH, Smittenaar P, Helms G, Lutti A, Weiskopf N. 2014. Widespread age-related differences in the human brain microstructure revealed by quantitative magnetic resonance imaging. Neurobiol Aging. 35:1862–1872.

Crossley NA, Mechelli A, Scott J, Carletti F, Fox PT, McGuire P, Bullmore ET. 2014. The hubs of the human connectome are generally implicated in the anatomy of brain disorders. Brain. 137:2382–2395.

Crossley NA, Mechelli A, Vertes PE, Winton-Brown TT, Patel AX, Ginestet CE, McGuire P, Bullmore ET. 2013. Cognitive relevance of the community structure of the human brain functional coactivation network. Proc Natl Acad Sci U S A. 110:11583–11588.

Dale AM, Fischl B, Sereno MI. 1999. Cortical surface-based analysis. I. Segmentation and surface reconstruction. Neuroimage. 9:179–194.

Deary IJ, Penke L, Johnson W. 2010. The neuroscience of human intelligence differences. Nat Rev Neurosci. 11:201–211.

Deoni SC, Dean DC, 3rd, Remer J, Dirks H, O’Muircheartaigh J. 2015. Cortical maturation and myelination in healthy toddlers and young children. Neuroimage. 115:147–161.

DiCiccio TJ, Efron B. 1996. Bootstrap confidence intervals. Stat Sci. 11:189–212.

Douaud G, Groves AR, Tamnes CK, Westlye LT, Duff EP, Engvig A, Walhovd KB, James A, Gass A, Monsch AU, Matthews PM, Fjell AM, Smith SM, Johansen-Berg H. 2014. A common brain network links development, aging, and vulnerability to disease. Proc Natl Acad Sci U S A. 111:17648–17653.

Evans AC. 2013. Networks of anatomical covariance. Neuroimage. 80:489–504.

Fjell AM, McEvoy L, Holland D, Dale AM, Walhovd KB, Alzheimer’s Disease Neuroimaging I. 2014. What is normal in normal aging? Effects of aging, amyloid and Alzheimer’s disease on the cerebral cortex and the hippocampus. Prog Neurobiol. 117:20–40.

Fjell AM, Walhovd KB, Westlye LT, Ostby Y, Tamnes CK, Jernigan TL, Gamst A, Dale AM. 2010. When does brain aging accelerate? Dangers of quadratic fits in cross-sectional studies. Neuroimage. 50:1376–1383.

Flechsig P. 1901. Developmental (myelogenetic) localisation of the cerebral cortex in the human subject. The Lancet. 158:1027–1030.

Gao W, Lin W, Chen Y, Gerig G, Smith JK, Jewells V, Gilmore JH. 2009. Temporal and spatial development of axonal maturation and myelination of white matter in the developing brain. AJNR Am J Neuroradiol. 30:290–296.

Gennari F. 1782. De peculiari structura cerebri nonnullisque ejus morbis … Paucae aliae anatom. observat. accedunt. In. Parma: Ex regio typographeo.

Glasser MF, Coalson TS, Robinson EC, Hacker CD, Harwell J, Yacoub E, Ugurbil K, Andersson J, Beckmann CF, Jenkinson M, Smith SM, Van Essen DC. 2016. A multi-modal parcellation of human cerebral cortex. Nature. 536:171–178.

Glasser MF, Goyal MS, Preuss TM, Raichle ME, Van Essen DC. 2014. Trends and properties of human cerebral cortex: correlations with cortical myelin content. Neuroimage. 93 Pt 2:165–175.

Glasser MF, Van Essen DC. 2011. Mapping human cortical areas in vivo based on myelin content as revealed by T1- and T2-weighted MRI. J Neurosci. 31:11597–11616.

Greve DN, Fischl B. 2009. Accurate and robust brain image alignment using boundary-based registration. Neuroimage. 48:63–72.

Grydeland H, Walhovd KB, Tamnes CK, Westlye LT, Fjell AM. 2013. Intracortical myelin links with performance variability across the human lifespan: results from T1- and T2-weighted MRI myelin mapping and diffusion tensor imaging. J Neurosci. 33:18618–18630.

Grydeland H, Westlye LT, Walhovd KB, Fjell AM. 2015. Intracortical Posterior Cingulate Myelin Content Relates to Error Processing: Results from T1- and T2-Weighted MRI Myelin Mapping and Electrophysiology in Healthy Adults. Cereb Cortex.

Harkins KD, Xu J, Dula AN, Li K, Valentine WM, Gochberg DF, Gore JC, Does MD. 2016. The microstructural correlates of T1 in white matter. Magn Reson Med. 75:1341–1345.

Hartigan JA, Hartigan P. 1985. The dip test of unimodality. The Annals of Statistics.70–84.

Herve PY, Zago L, Petit L, Mazoyer B, Tzourio-Mazoyer N. 2013. Revisiting human hemispheric specialization with neuroimaging. Trends Cogn Sci. 17:69–80.

Huntenburg JM, Bazin PL, Goulas A, Tardif CL, Villringer A, Margulies DS. 2017. A Systematic Relationship Between Functional Connectivity and Intracortical Myelin in the Human Cerebral Cortex. Cereb Cortex. 27:981–997.

Kabaso D, Coskren PJ, Henry BI, Hof PR, Wearne SL. 2009. The electrotonic structure of pyramidal neurons contributing to prefrontal cortical circuits in macaque monkeys is significantly altered in aging. Cereb Cortex. 19:2248–2268.

Kail R. 1991. Developmental change in speed of processing during childhood and adolescence. Psychol Bull. 109:490–501.

Kessaris N, Fogarty M, Iannarelli P, Grist M, Wegner M, Richardson WD. 2006. Competing waves of oligodendrocytes in the forebrain and postnatal elimination of an embryonic lineage. Nat Neurosci. 9:173–179.

Kochunov P, Williamson DE, Lancaster J, Fox P, Cornell J, Blangero J, Glahn DC. 2012. Fractional anisotropy of water diffusion in cerebral white matter across the lifespan. Neurobiol Aging. 33:9–20.

Lerch JP, Worsley K, Shaw WP, Greenstein DK, Lenroot RK, Giedd J, Evans AC. 2006. Mapping anatomical correlations across cerebral cortex (MACACC) using cortical thickness from MRI. Neuroimage. 31:993–1003.

Lindenberger U, von Oertzen T, Ghisletta P, Hertzog C. 2011. Cross-sectional age variance extraction: what’s change got to do with it? Psychol Aging. 26:34–47.

Lopez-Otin C, Blasco MA, Partridge L, Serrano M, Kroemer G. 2013. The hallmarks of aging. Cell. 153:1194–1217.

Marra G, Wood SN. 2012. Coverage properties of confidence intervals for generalized additive model components. Scandinavian Journal of Statistics. 39:53–74.

Miller DJ, Duka T, Stimpson CD, Schapiro SJ, Baze WB, McArthur MJ, Fobbs AJ, Sousa AM, Sestan N, Wildman DE, Lipovich L, Kuzawa CW, Hof PR, Sherwood CC. 2012. Prolonged myelination in human neocortical evolution. Proc Natl Acad Sci U S A. 109:16480–16485.

Paus T, Keshavan M, Giedd JN. 2008. Why do many psychiatric disorders emerge during adolescence? Nat Rev Neurosci. 9:947–957.

Paus T, Zijdenbos A, Worsley K, Collins DL, Blumenthal J, Giedd JN, Rapoport JL, Evans AC. 1999. Structural maturation of neural pathways in children and adolescents: in vivo study. Science. 283:1908–1911.

Peters A. 2009. The effects of normal aging on myelinated nerve fibers in monkey central nervous system. Front Neuroanat. 3:11.

Rakic P. 2002. Neurogenesis in adult primate neocortex: an evaluation of the evidence. Nat Rev Neurosci. 3:65–71.

Raz N. 1999. Aging of the brain and its impact on cognitive performance: Integration of structural and functional findings. In. The handbook of aging and cognition 2nd ed. Mahwah, N.J: Lawrence Erlbaum.

Raz N, Gunning-Dixon F, Head D, Rodrigue KM, Williamson A, Acker JD. 2004. Aging, sexual dimorphism, and hemispheric asymmetry of the cerebral cortex: replicability of regional differences in volume. Neurobiol Aging. 25:377–396.

Righart R, Biberacher V, Jonkman LE, Klaver R, Schmidt P, Buck D, Berthele A, Kirschke JS, Zimmer C, Hemmer B, Geurts JJG, Muhlau M. 2017. Cortical pathology in multiple sclerosis detected by the T1/T2-weighted ratio from routine magnetic resonance imaging. Ann Neurol. 82:519–529.

Ritchie J, Pantazatos SP, French L. 2018. Transcriptomic characterization of MRI contrast with focus on the T1-w/T2-w ratio in the cerebral cortex. Neuroimage. 174:504–517.

Roalf DR, Quarmley M, Elliott MA, Satterthwaite TD, Vandekar SN, Ruparel K, Gennatas ED, Calkins ME, Moore TM, Hopson R, Prabhakaran K, Jackson CT, Verma R, Hakonarson H, Gur RC, Gur RE. 2016. The impact of quality assurance assessment on diffusion tensor imaging outcomes in a large-scale population-based cohort. Neuroimage. 125:903–919.

Rosen AFG, Roalf DR, Ruparel K, Blake J, Seelaus K, Villa LP, Ciric R, Cook PA, Davatzikos C, Elliott MA, Garcia de La Garza A, Gennatas ED, Quarmley M, Schmitt JE, Shinohara RT, Tisdall MD, Craddock RC, Gur RE, Gur RC, Satterthwaite TD. 2018. Quantitative assessment of structural image quality. Neuroimage. 169:407–418.

Rowitch DH, Kriegstein AR. 2010. Developmental genetics of vertebrate glial-cell specification. Nature. 468:214–222.

Rubinov M, Sporns O. 2010. Complex network measures of brain connectivity: uses and interpretations. Neuroimage. 52:1059–1069.

Salthouse TA. 1996. The processing-speed theory of adult age differences in cognition. Psychol Rev. 103:403–428.

Satterthwaite TD, Wolf DH, Loughead J, Ruparel K, Elliott MA, Hakonarson H, Gur RC, Gur RE. 2012. Impact of in-scanner head motion on multiple measures of functional connectivity: relevance for studies of neurodevelopment in youth. Neuroimage. 60:623–632.

Sawiak SJ, Shiba Y, Oikonomidis L, Windle CP, Santangelo AM, Grydeland H, Cockcroft G, Bullmore ET, Roberts AC. 2018. Trajectories and Milestones of Cortical and Subcortical Development of the Marmoset Brain From Infancy to Adulthood. Cereb Cortex.

Scholtens LH, de Reus MA, de Lange SC, Schmidt R, van den Heuvel MP. 2016. An MRI Von Economo - Koskinas atlas. Neuroimage.

Seldon HL. 2005. Does brain white matter growth expand the cortex like a balloon? Hypothesis and consequences. Laterality. 10:81–95.

Shafee R, Buckner RL, Fischl B. 2015. Gray matter myelination of 1555 human brains using partial volume corrected MRI images. Neuroimage. 105:473–485.

Shafto MA, Tyler LK. 2014. Language in the aging brain: the network dynamics of cognitive decline and preservation. Science. 346:583–587.

Shinn M, Romero-Garcia R, Seidlitz J, Vasa F, Vertes PE, Bullmore E. 2017. Versatility of nodal affiliation to communities. Sci Rep. 7:4273.

Sigalovsky IS, Fischl B, Melcher JR. 2006. Mapping an intrinsic MR property of gray matter in auditory cortex of living humans: a possible marker for primary cortex and hemispheric differences. Neuroimage. 32:1524–1537.

Solari SV, Stoner R. 2011. Cognitive consilience: primate non-primary neuroanatomical circuits underlying cognition. Front Neuroanat. 5:65.

Sporns O, Betzel RF. 2016. Modular Brain Networks. Annu Rev Psychol. 67:613–640.

Stuber C, Morawski M, Schafer A, Labadie C, Wahnert M, Leuze C, Streicher M, Barapatre N, Reimann K, Geyer S, Spemann D, Turner R. 2014. Myelin and iron concentration in the human brain: a quantitative study of MRI contrast. Neuroimage. 93 Pt 1:95–106.

Tau GZ, Peterson BS. 2010. Normal development of brain circuits. Neuropsychopharmacology : official publication of the American College of Neuropsychopharmacology. 35:147–168.

Triarhou LC. 2007. The Economo-Koskinas atlas revisited: cytoarchitectonics and functional context. Stereotact Funct Neurosurg. 85:195–203.

van den Heuvel MP, Sporns O. 2013. Network hubs in the human brain. Trends Cogn Sci. 17:683–696.

van der Knaap MS, Valk J. 1990. MR imaging of the various stages of normal myelination during the first year of life. Neuroradiology. 31:459–470.

Vertes PE, Rittman T, Whitaker KJ, Romero-Garcia R, Vasa F, Kitzbichler MG, Wagstyl K, Fonagy P, Dolan RJ, Jones PB, Goodyer IM, Consortium N, Bullmore ET. 2016. Gene transcription profiles associated with inter-modular hubs and connection distance in human functional magnetic resonance imaging networks. Philos Trans R Soc Lond B Biol Sci. 371.

Vincent JL, Patel GH, Fox MD, Snyder AZ, Baker JT, Van Essen DC, Zempel JM, Snyder LH, Corbetta M, Raichle ME. 2007. Intrinsic functional architecture in the anaesthetized monkey brain. Nature. 447:83–86.

Walhovd KB, Fjell AM, Giedd J, Dale AM, Brown TT. 2017. Through Thick and Thin: a Need to Reconcile Contradictory Results on Trajectories in Human Cortical Development. Cereb Cortex. 27:1472–1481.

Westlye LT, Walhovd KB, Dale AM, Bjornerud A, Due-Tonnessen P, Engvig A, Grydeland H, Tamnes CK, Ostby Y, Fjell AM. 2010. Differentiating maturational and aging-related changes of the cerebral cortex by use of thickness and signal intensity. Neuroimage. 52:172–185.

Westlye LT, Walhovd KB, Dale AM, Espeseth T, Reinvang I, Raz N, Agartz I, Greve DN, Fischl B, Fjell AM. 2009. Increased sensitivity to effects of normal aging and Alzheimer’s disease on cortical thickness by adjustment for local variability in gray/white contrast: a multi-sample MRI study. Neuroimage. 47:1545–1557.

Whitaker KJ, Vertes PE, Romero-Garcia R, Vasa F, Moutoussis M, Prabhu G, Weiskopf N, Callaghan MF, Wagstyl K, Rittman T, Tait R, Ooi C, Suckling J, Inkster B, Fonagy P, Dolan RJ, Jones PB, Goodyer IM, Consortium N, Bullmore ET. 2016. Adolescence is associated with genomically patterned consolidation of the hubs of the human brain connectome. Proc Natl Acad Sci U S A. 113:9105–9110.

Wood SN. 2006. Generalized Additive Models: An Introduction with R. Boca Raton, FL: CRC Press.

Yakovlev PI, Lecours A-R. 1967. The myelogenic cycles of regional maturation of the brain. In: Minkowski A, editor. Regional Development of the Brain in Early Life Oxford and Edinburg: Blackwell Scientific Publications p 3–70.

Yarkoni T, Poldrack RA, Nichols TE, Van Essen DC, Wager TD. 2011. Large-scale automated synthesis of human functional neuroimaging data. Nat Methods. 8:665–670.

Yeatman JD, Wandell BA, Mezer AA. 2014. Lifespan maturation and degeneration of human brain white matter. Nat Commun. 5:4932.

Zalc B, Colman DR. 2000. Origins of vertebrate success. Science. 288:271–272.

